# Molecular Dynamics and Water site bias docking method allows the identification of key amino acids in the Carbohydrate Recognition Domain of a viral protein

**DOI:** 10.1101/2023.06.01.543333

**Authors:** Marcelo D. Gamarra, Maria Eugenia Dieterle, Juan I. Blanco Capurro, Leandro Radusky, Mariana Piuri, Carlos P. Modenutti

**Affiliations:** Departamento de Química Biológica, Facultad de Ciencias Exactas y Naturales, Universidad de Buenos Aires (FCEyN-UBA) e Instituto de Química Biológica de la Facultad de Ciencias Exactas y Naturales (IQUIBICEN) CONICET, Pabellón 2 de Ciudad Universitaria, Ciudad de Buenos Aires C1428EHA, Argentina

## Abstract

Carbohydrate-binding modules (CBMs) are protein domains that typically reside near catalytic domains, increasing substrate-protein proximity by constraining the conformational space of carbohydrates. Due to the flexibility and variability of glycans, the molecular details of how these protein regions recognize their target molecules are not always fully understood. Computational methods, including molecular docking and molecular dynamics simulations, have been employed to investigate lectin-carbohydrate interactions. In this study, we introduce a novel approach that integrates multiple computational techniques to identify the critical amino acids involved in the interaction between a CBM located at the tip of bacteriophage J-1’s tail and its carbohydrate counterparts. Our results highlight three amino acids that play a significant role in binding, which we confirmed through in vitro experiments. By presenting this approach, we offer an intriguing alternative for pinpointing amino acids that contribute to protein-sugar interactions, leading to a more thorough comprehension of the molecular determinants of lectin-carbohydrate interactions.

## Introduction

Lectins are a large group of biomolecules involved in many biological processes, including cell recognition, communication, and cell growth (Vijayan and Chandra 1999). They participate in recognizing and eliminating pathogens as well as facilitating the entry of microbial pathogens and parasites into the host (Baum et al. 2014; Lujan et al. 2018). All these mechanisms are directly related to the structure of lectins and their ability to selectively recognize different patterns in carbohydrate polymers (C. Modenutti et al. 2015). Knowing how the lectin-carbohydrate interaction occurs is critical to understanding the biological processes and developing strategies for infection control.

Bacteriophages are viruses that specifically infect bacteria. At the tip of the tail, a phage-dedicated host recognition machinery interacts with different components of the bacterial envelope (Dunne et al. 2019). J-1 phage (a Siphovirus) is a well-known bacteriophage of great importance in the dairy industry because it can infect several *Lactobacillus casei* / *paracase*i strains (Sechaud et al. 1988; Capra et al. 2006; M. E. Dieterle et al. 2014a). Although the recognition machinery of many viruses is known, the molecular mechanism by which the J-1 phage recognizes its host has not been completely revealed (Veesler and Cambillau 2011; Spinelli et al. 2014). In addition to its structural role in baseplate formation, the J-1 distal tail protein (Dit) is essential for recognizing *L. casei* receptors located in the cell envelope (M. E. Dieterle et al. 2014b). In J-1 phage, Dit protein has two carbohydrate-binding modules (CBM1 and CBM2), but only CBM2 seems to be involved in bacterial cell wall polysaccharides (CWPs) recognition (M. E. Dieterle et al. 2014b; Vinogradov et al. 2016a; M.-E. Dieterle et al. 2017a). The CWP of *L. casei* BL23 is rich in a-Rhamnose (Vinogradov et al. 2016b) and previous results showed that J-1 phage adsorption to *L. casei* cells was inhibited in the presence of this monosaccharide, leading to the idea that the interaction must be mediated by rhamnose (Yokokura 1971; M. E. Dieterle et al. 2014b; M.-E. Dieterle et al. 2017b). In 2017, the first structure of J-1 CBM2 was deposited at the Protein Data Bank (PDB) and showed a folding similar to lectins (PDB ID 5LY8). Despite the superb resolution of the CBM2 module in its apo form (1.28 Å), a structure of the complex with rhamnose has not been obtained yet. This is likely because the carbohydrate-binding site undergoes a conformational change upon ligand binding, leading to crystal destabilization. Through a comparative analysis of the CBM2 structure with other lectins, Dieterle et al. were able to propose a putative binding site and potential interaction with monosaccharides. However, to fully comprehend the molecular determinants of the interaction with CWPs, it is necessary to study a protein-ligand complex that contains at least one oligosaccharide.

X-ray crystallography is one of the most popular methods used to obtain a detailed description of protein-ligand interactions. However, solving the three-dimensional structure of carbohydrates is a well-known challenge because of the inherent flexibility of glycans (Nagae and Yamaguchi 2012; Gabius et al. 2011). Bioinformatics methods are a powerful alternative for structure modeling. Different methods have been developed for carbohydrate docking as BALLDock/SLICK (Kerzmann et al. 2008), Vina-Carb (Nivedha et al. 2016), and Glycotroch-Vina (E. Boittier et al. 2020) showing acceptable performance for protein-carbohydrates structure prediction. However, several challenges are still unsolved for oligomers larger than trisaccharides, especially those concerning flexibility and the twists they can take (Xiong et al. 2015). Identifying regions with high probability of finding water molecules, the so-called Water Sites (WS), and incorporating it to the docking protocol can lead to more accurate carbohydrate docking prediction (Gauto et al. 2013; C. Modenutti et al. 2015; Blanco Capurro et al. 2019). This is because the -OH groups of carbohydrates tend to mimic water interactions and form similar hydrogen bonding networks when making contact with protein surfaces (Saraboji et al. 2012; Gauto et al. 2009, 2011; Di Lella et al. 2007). Consequently, it is crucial to characterize the solvent structure at the carbohydrate-binding site (CBS), and incorporating this information to modify the scoring function for docking can enhance the performance of docking simulations.

Here, we developed a pipeline that contributes to the understanding of the binding mode of phage J-1 with its ligand and identifies key amino acids involved in the carbohydrate recognition domain.

## Results

The results are organized as follows: initially, different methods were combined to identify the carbohydrate-binding site of CBM2 and the amino acids involved in protein-ligand interactions. Our approach (Figure 1) involved both quantitative and qualitative estimation of the contribution of the receptor amino acids to the interaction with the solvent and the ligands. Applying these methods allowed us to reach a consensus on the molecular determinants of the interaction. Subsequently, we validated the most promising candidates by estimating the impact on the protein-ligand interaction using FoldX and molecular dynamics simulations. Finally, we confirmed the significance of the proposed amino acids by conducting *in vitro* assays to test the binding between the bacteria and the protein.

**Figure 1:**
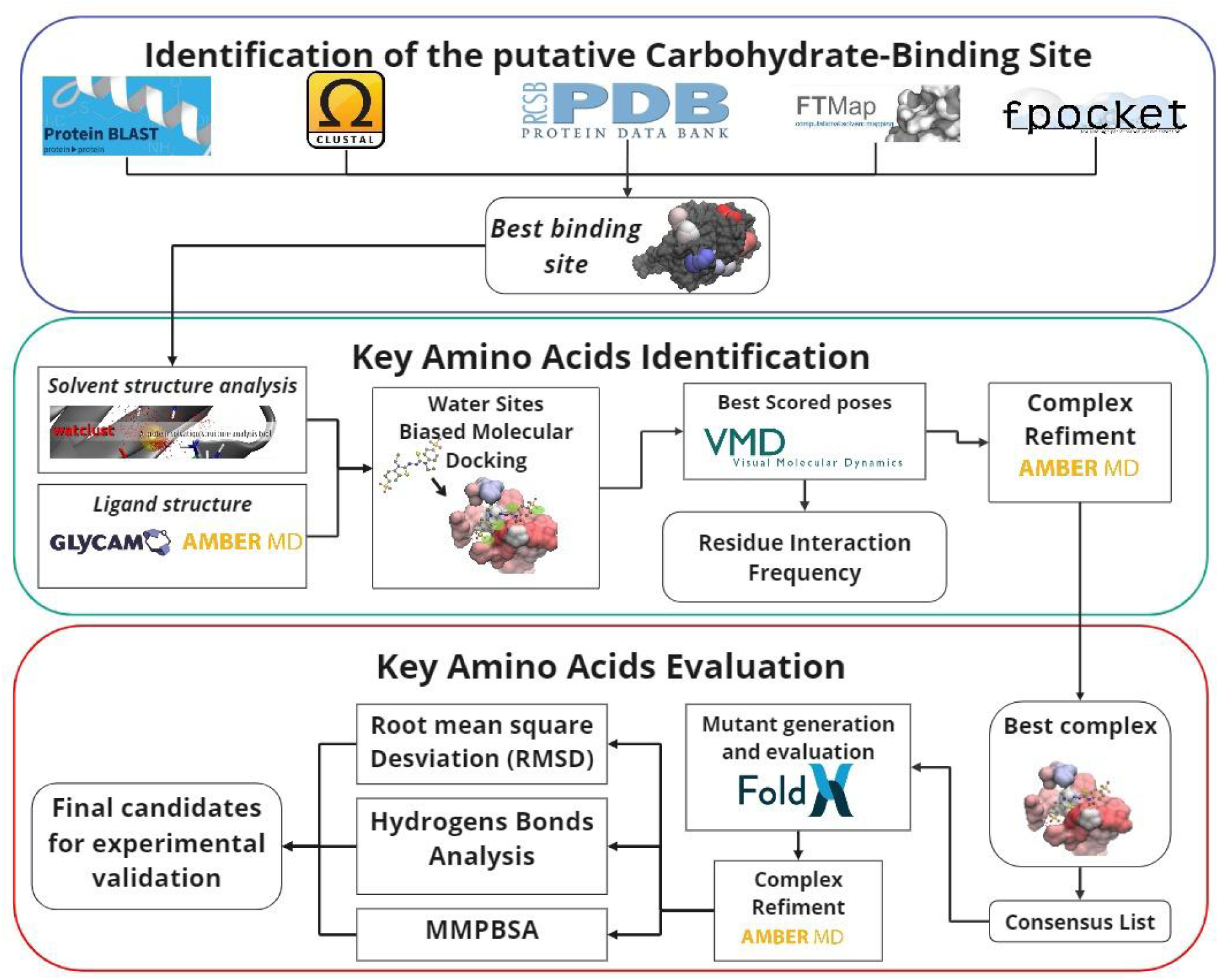
Bioinformatic approach for key amino acid identification and evaluation in the CBM2 of bacteriophage J-1’s tail. The analysis started with the binding site identification, combining sequence (BLASTP and ClustalO) and structure (structure and motif PDB Search) similarity search, pockets identification (Fpocket), and ligand hotspot mapping (FTMap). For the critical amino acids detection we used molecular dynamics simulations (AMBER) followed by water sites biased docking (AutoDock Bias). Finally, the evaluation of the candidates was performed by estimating the stability of the complex (RMSD, hydrogen bond frequency, and interaction energy) using the AMBER package.

In the second stage of the study, a quantitative dynamic characterization was conducted. This involved simulating molecular dynamics of previously obtained complexes and measuring polar interactions through hydrogen bonds and MM-GBSA parameters. Additionally, *in silico* mutation analysis was performed using FoldX to evaluate the impact of mutations in key amino acids on ligand interaction stability (Schymkowitz et al. 2005).

### Identification of the putative Carbohydrate-Binding Site of CBM2

#### Protein sequence mining and analysis

The analysis started by identifying the potential binding site through which CBM2 recognizes and attaches to carbohydrates in the host’s wall. Two bioinformatics approaches were employed for its detection. Firstly, assuming that regions important for protein-ligand interaction should be conserved a sequence identity analysis was conducted by searching information in databases. Using the Uniref50 database (https://www.uniprot.org/) we retrieved 20 protein sequences identified as distal tail proteins (DITs) from various lactic acid bacteria phages and performed a multiple sequence alignment (MSA) (Table S1). The analysis revealed sequence identities ranging from 85% to 95% for the CBM2 domain, and in many cases even higher.

Because the high identity between sequences difficult the identification of conserved regions, a more exhaustive search was carried out by aligning the sequences against the RefSeq database (https://www.ncbi.nlm.nih.gov/refseq/) (Figure S1). While the profile exhibited low identity (ranging between 22% to 50%) among the protein sequences, certain positions showed significant conservation. Interestingly, half of the conserved amino acids were located far apart in sequence (except the WXGP motif, corresponding to W440, G442, and P443 in CBM2 Uniprot numbering) but closely situated in the three-dimensional structure of CBM2, with all of them located at a distance ≤ 5Å from W440 (Figure 2). Analyzing the CBM2 structure (PDB ID 5LY8) in more detail, we found that all the side chains of the MSA-conserved amino acids were buried within the protein’s core, except for W440, which was partially exposed to the solvent. These findings led us to hypothesize that the WXGP motif could be involved in carbohydrate interaction but the searches in the PROSITE database returned no results. In order to probe our hypothesis about the WXGP motif, we performed a pattern search against the Protein Data Bank (PDB), restricting the search only to those proteins co-crystallized with carbohydrates. The search results revealed that most of the identified proteins present an interaction between carbohydrates and the WXGP motif (Figure 2-B and Table S3).

**Figure 2.**
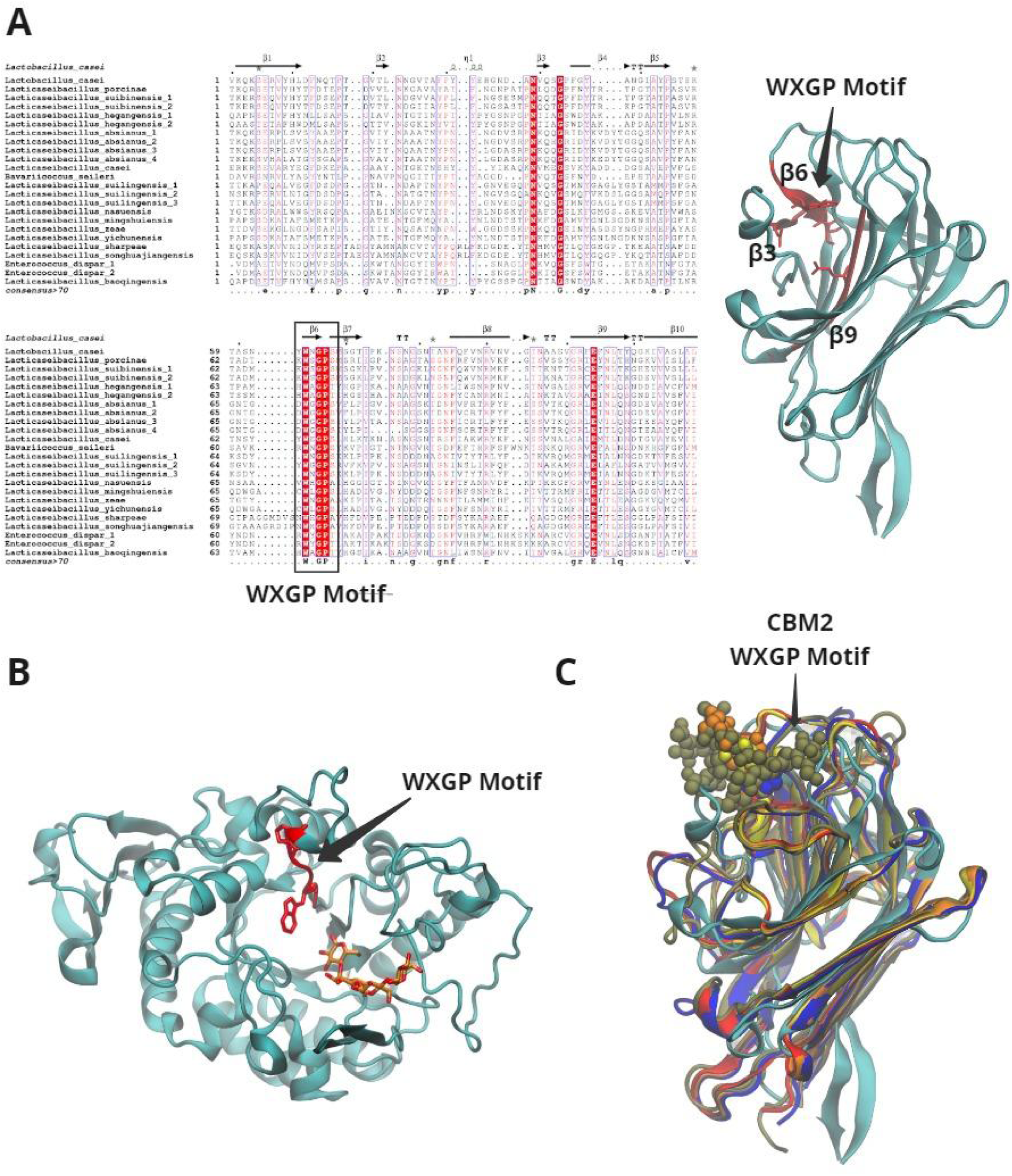
Sequence and structure similarity search analysis over Carbohydrate Bindign Module 2. **A.** Multiple alignments of the CBM2 sequence with similar proteins found with *RefSeq*, on the right conserved amino acids throughout the alignment represented on the three-dimensional structure of CBM2. **B.** Carbohydrate-binding protein with the WXGP motif present at the binding site (PDB ID 1JDC). **C.** Comparison of proteins with CBM2 similar folding (PDBIDs: 5LY8, 2E51, 5AVA, 2E7T, 1FAY, 1WBL), found through the search carried out with PDB (Protein Data Bank), The carbohydrates attached to each structure are represented with spheres.

Although these findings support the hypothesis that the WXGP motif was involved in ligand binding, the folding of the proteins retrieved was very different. Based on the concept that proteins exhibiting similar folding patterns may also have similar ligand-binding mechanisms, a similarity folding search was performed against the PDB using CBM2 structure as a query. Special attention was given to structures with co-crystallized ligands, as their positioning could potentially provide insights into potential binding sites. The majority of the proteins found were of the lectin type (Table S4). The structural alignment of these proteins revealed that the ligand-binding sites correlated with the region where the conserved amino acids detected in the sequence alignment were located, reinforcing the idea of a possible binding site (Figure 2-C). All these findings suggest a potential binding site centered around the conserved amino acids W440.

#### Cavities and ligand hot sports identification and characterization

Next, we wanted to identify the areas at the protein surface with characteristics of ligand binding sites as an independent validation of the previously identified region. For this purpose first, we used FPocket software (Le Guilloux, Schmidtke, and Tuffery 2009) and mapped all the possible cavities over the CBM2 domain and sugar-binding proteins with similar folding (Figure 3-A,B).

**Figure 3:**
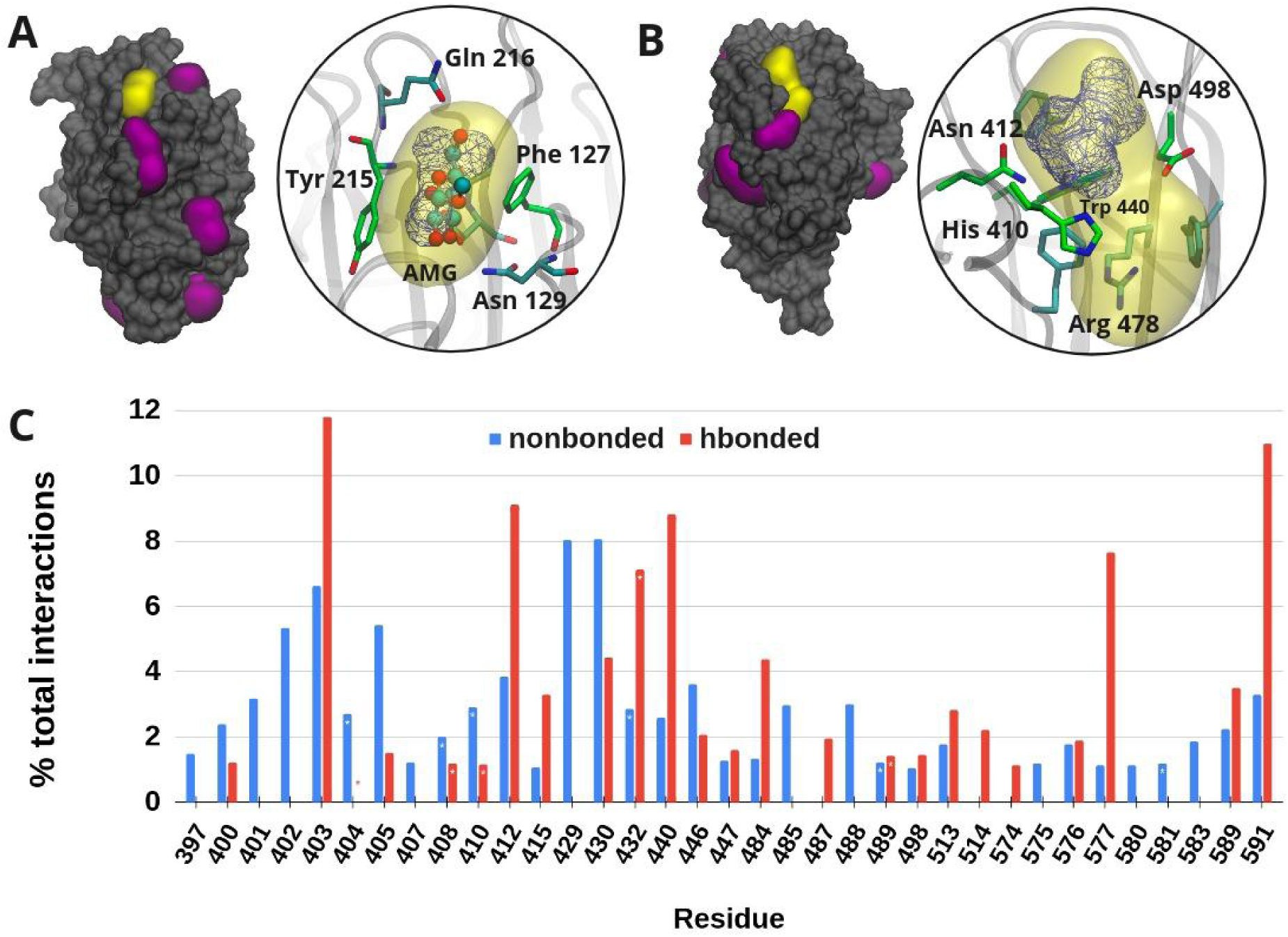
Cavities identification and characterization in CBM2 and proteins with similar folding. **A.** Representation of winged bean acidic lectin complexed with methyl-alpha-d-galactose (AMG) (PDB ID: 1FAY). **B.** The Carbohydrate binding module 2 (PDB ID: 5LY8). The cavities detected by Fpocket are shown in purple and putative pocket binding sites detected are shown in yellow while high-density regions occupied by FTMap probes are shown as blue lines. The circles beside panels A and B show the potential key amino acids present in the active site detected. The carbon atoms of the amino acids identified with FTMap are shown in green, while the amino acids within a distance of 3.5 Å from the alpha spheres of FPocket are shown in cyan. In the case of 1FAY, the ligand AMG is represented with ball and stick and colored by atom type. **C.** Percentage of interaction for polar and nonpolar contacts between FTMap probes and CBM2 protein residues. The asterisks in the columns highlight the amino acids belonging to the putative binding site.

FPocket was applied to a protein dataset consisting of structures with a ligand bound and a folding pattern similar to CBM2. Prior to analysis, the ligand and water molecules were removed from the protein surface with the aim to assess the sensitivity of FPocket in detecting cavities onto sugar binding proteins (specifically if the algorithm was able to identify the carbohydrate binding sites). FPocket successfully identified the ligand binding sites in all cases, along with additional pockets of interest (Figure 3-A). Notably, when applied to CBM2, FPocket revealed a significant pocket surrounding the conserved amino acids W440 among the results (Figure 3-B). As anticipated for lectin-like proteins, where the cavities are typically not well defined, the druggability scores (DS) were below 0.5 for the analyzed structures, unlike conventional drug-binding sites that typically exhibit DS values above this threshold (L. Radusky et al. 2014). Despite this, the best-scored cavity in CBM2 (DS 0.46) was located adjacent to the W440 of the WXGP motif, including the residues F405, H410, N412, R478, F496, D498, and F583.

Finally, we evaluated the ability of the potential CBS to interact with ligands, particularly carbohydrates. To achieve this objective, the FTMap server was employed, which utilizes a diverse range of small organic probe molecules possessing various physicochemical properties (Ngan et al. 2012). This server identifies surface regions on macromolecules that significantly contribute to ligand binding free energy or act as binding hot spots.

When using FTMap, similar results to FPocket were obtained, indicating a correlation between cavities presence and the ability to interact with small ligands. Again, we used winged bean acidic lectin complexed with methyl-alpha-d-galactose (AMG) as a positive control for our analysis. FTMap successfully detected a hotspot at the binding site of the ligand AMG, as observed in the FPocket results (Figure 3-A). Regarding CBM2, the findings indicate the presence of multiple hotspots on the surface of CBM2, with one of them aligning with the previously identified cavity adjacent to the W440 residue (Figure 3, panel B). Notably, the majority of the probes clustered in this hotspot exhibited similar chemical properties to sugars, such as ethanol, isobutanol, and acetaldehyde, among others. Charged and aromatic molecules did not cluster in this area, and the primary interactions observed were polar, involving the sidechain of residues H410, N412, W440, and D498 (Figure 3-C). These results suggest the potential involvement of these residues in the initial recognition of carbohydrates. Combining both approaches, we concluded that the carbohydrate-binding site (CBS) of CBM2 is likely located around W440, prompting further analysis of this region.

### Identification of critical amino acid carbohydrate binding

#### Carbohydrate binding site characterization by molecular dynamics simulations

Solvent structure in the carbohydrate-binding site (CBS) tends to mimic the carbohydrate hydroxyl positions in the protein-sugar complexes, and thus the identification of the Water Sites (WS) allows us to predict possible protein-ligand interactions. For a detailed characterization of the previous identify CBS, we carried out a solvent structure analysis, which has proven that it is a powerful tool for identifying potential polar contacts between protein and carbohydrates <u>(Gauto et al. 2013; Modenutti et al. 2015)</u>. First, using explicit solvent MD, we calculated the water sites (WS) around the same proposed binding site (see methods). Next, the so-called “crystallographic waters sites (CWS)” around the putative binding site of CBM2 (PDBID: 5LY8) were analyzed. To identify the main protein interactions, those WS that corresponded to CWS were considered interaction reporter sites, i.e. all the WS that were within 1.4A of a CWS were considered as the same site. As a result, residues N412, W440, and D498 were identified as the amino acids with the highest frequency of interaction with the solvent.

##### Water site bias molecular docking

Due to the high number of probable conformations that CBM2 ligands polysaccharides could adopt, docking experiments were followed. To reveal the ligand binding mode and to determine the amino acids that interact more frequently around all docking results. We previously showed the MD ability to sample ‘holo-like’ conformational structures. In this sense, the CBM2 conformational diversity was considered by obtaining an ensemble of receptor conformations derived from MD snapshots. All three ligand variants of CWPS were used for docking assays. They were prepared in a method that allows us to reduce the variability of conformations using explicit waters MD and a clustering algorithm to obtain the most energy-stable conformations for each ligand, with the correct Phi and Psi angles (Figure 4-C, See Methods).

**Figure 4.**
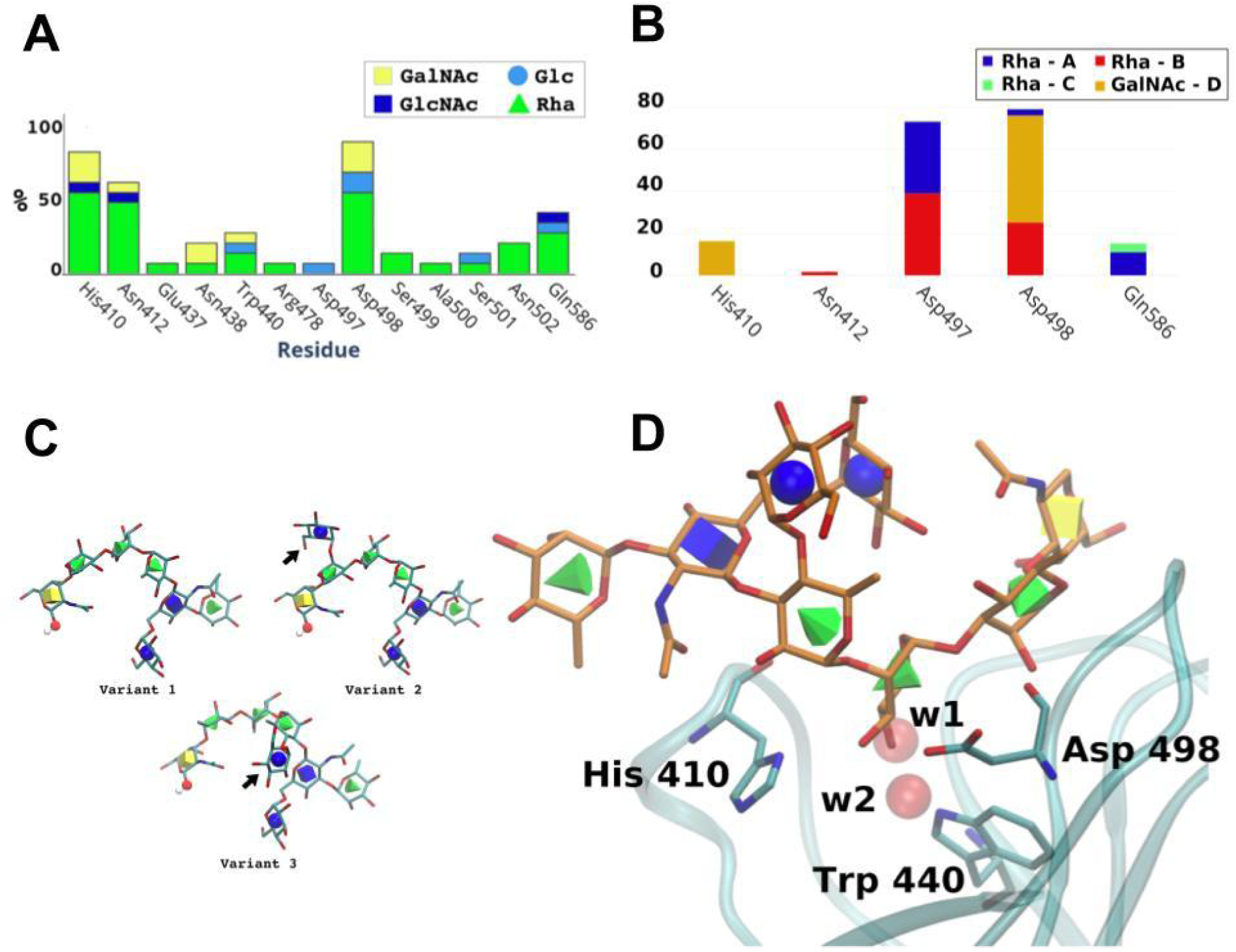
Identification of relevant amino acids in the CBM2 binding site. **A.** Frequency histogram of the residue interactions showed in the best-ranked docking results. **B.** Hydrogen bonds frequency of interaction for each CBS residue within molecular dynamics simulations. **C.** Cell wall polysaccharide variants built with Glycam server. Variants are similar but only differ in a Glucose substituent indicated with arrows. GalNAc’s anomeric hydroxyl group of the reducing end is highlighted as balls. **D.** Possible binding mode of CWPS variant 3, showing the Rhamnose interaction with residues H410, W440, and D497. All geometric shapes and colors correspond to the standard SNFG code for carbohydrates (Thieker et al. 2016). w1 and w2 water sites considered for WSBDM correspond to 443 and 496 crystal waters respectively.

For molecular docking simulations, the identify WS were used to modify the Autodock4 scoring function, and the *solvent site bias docking method* (SSBDM) was performed <u>(Arcon et al. 2019)</u>. Each receptor conformation from the ensemble was used for docking ligands, and all docking results were combined. Polar interaction analysis was realized for the selected poses by measuring the frequency of interaction for each CBS residue. The results showed that the amino acids with the highest frequency of interaction were H410, N412, Q586, and D498 (Figure 4-A). In addition, it was observed that rhamnose residues perform the most frequent interaction at the CBS, although other moieties such as GlcNAc or Glc may occur less frequently.

#### Ligand stability evaluation

The objective of this first part was two-fold: firstly, to identify the key amino acids involved in the interaction between CBM2 protein and polysaccharides, and secondly, to determine the most likely mode of binding or recognition between them. A stable protein-ligand complex is formed during the binding process, and molecular dynamics can be used to assess the stability of the complex over time. Therefore, for each of the best results obtained from the docking experiments between CBM2 and each variant of the CWPs, we performed 10ns of MD simulations to evaluate the stability of the complex. Our results showed that variant 3 (Figure 4 - C,D) was the most stable complex, and this was used for subsequent evaluations.

#### Protein stability evaluation

Finally, Using the pyFoldX library, we identified amino acids crucial for the CBM2-polysaccharide interaction, while also considering their impact on protein stability when mutated. We found that H410, N412, Q586, and D498 were important for the interaction, but when we mutated W440, the protein folding became unstable, indicating that W440 is crucial for both the interaction and protein stability. Thus, W440 was considered key for the interaction, but we excluded it from subsequent steps of our proposed pipeline because of its destabilizing effect on protein folding (Table SI4).

### Validation of the relevant amino acids of the putative CBM2 binding site

#### Key Amino Acids Evaluation by explicit water molecular dynamics

The CBM2-carbohydrate complexes were evaluated through MD simulation to gain a deeper understanding of protein-ligand interactions. *Hydrogen bond analysis* was performed on the resulting trajectory to estimate the contribution of individual amino acids in the putative binding site (Figure 4 - B). The results revealed that the most frequent interactions occur through the D498 and D497 residues, with the latter playing a crucial role in protein-ligand interaction despite being overlooked in previous docking experiments due to its orientation within the protein crystal and the receptor’s flexibility. These results demonstrate the ability of MD to uncover important amino acids that may not have been identified through docking experiments alone.

#### *In silico* mutants evaluation

The complex selected as the most probable of the docking results was tested using FoldX. FoldX allows testing on the one hand the stability of an apo protein (without ligand) and on the other the stability of the protein-carbohydrate complex as a result of point mutations in the receptor. The amino acids used for replacement were selected based on the prediction by FoldX of minimal disturbance of the protein folding (Table SI4) (Schymkowitz et al. 2005). In relation to the protein-carbohydrate complex, causing point mutations destabilizes the binding of the ligand, being the double and triple mutant (D497L-D498L.H410F) which generates the greatest disturbance in the interaction (Table 1).

**Table 1.** Prediction of ligand stability energy in CBM2 and interaction energy stability of CBM2 - variant 3 ligand as a result of point mutations.

| Variant | $\Delta\Delta G$ (FoldX) | $\Delta\Delta G$ (MM-PBSA) |
| --- | --- | --- |
| H410F | +2.74 | +3.59 |
| D497L | +0.71 | +0.21 |
| D498L | +2.56 | +1.79 |
| D497L - D498L | +4.51 | +1.90 |
| D497L-D498L-H410F | +5.97 | +10.99 |

For the residues previously identified, those that were most representative in the analyzes were considered for further experiments. Specific *in silico* mutants were built and apo-protein stability and protein-ligand interaction were evaluated. Three different mutants were selected for analysis: the single mutant H410F, the double D497L-D498L, and the triple mutant H410F-D498L-D498L. MD simulations were performed for the three mutants to determine the stability of the protein-ligand complex and possible loss of the interaction (Makeneni, Thieker, and Woods 2018).

All mutants showed less stability compared to the *wild-type (WT)* based on the estimation of the Root Mean Square Deviation (RMSD) (**Figure 5-B**). The H410F and H410F-D498L-D498L mutants were the most affected in the ligand binding. This fact was due to the loss of donor/acceptor interactions caused by the adopted changes, which prompted us to evaluate the MD-ligand interaction energy with MM-GBSA (**Figure 5-C**). MM-GBSA Total showed that the interaction energy decreased in the complex as the number of mutations increased, with the triple mutant D497L being the most damaged, a result similar to the estimate made with FoldX (Table 1). The results showed that all mutants affect binding energy and the highest delta ΔG variations (ΔΔG) were seen in Rha-A, Rha-B and Rha-C (**Figure 5-A**).

**Figure 5.**
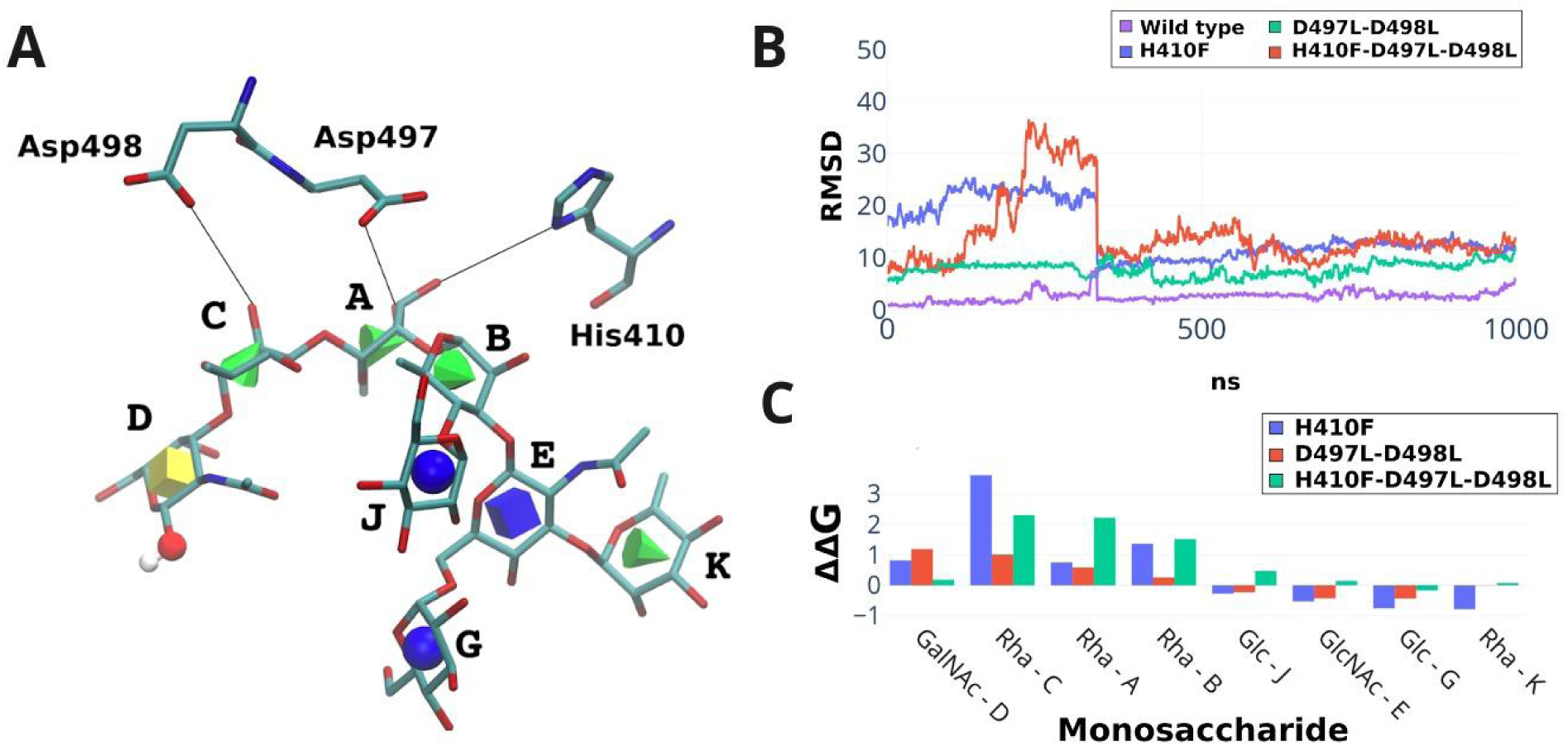
Validation of the relevant amino acids of the putative CBM2 binding site. **A.** Relevant amino acids in the CBM2-carbohydrate interaction. The main interaction occurs with the rhamnose monosaccharide (green cones). **B.** Stability of the CBM2-carbohydrate interaction in Molecular Dynamics simulations measured by RMSD. **C.** MD-ligand interaction energy estimation by MM-GBSA.

As expected, many amino acids were involved in the recognition of CWPS by CBM2, although only certain monosaccharides of the ligand interact more frequently (**Figure 4A, C**). The frequent interactions occur by the D497, D498 and less frequently by H410 residues (**Figure 4-B**) and point mutations in these amino acids cause less ligand stability at the binding site (**Figure 5-B, C**).

#### Effect of point mutations in CBM2 domain binding to host cells

To evaluate experimentally the role of the residues identified through bioinformatic analysis in the CBS of CBM2, we constructed a chimeric green fluorescent protein in frame with CBM2 (GFP-CBM2) protein and the derivative mutants. The recombinant proteins were tested using a binding assay to *L. casei* BL23 cells as previously described (Makeneni, Thieker, and Woods 2018; M. E. Dieterle et al. 2014a; M.-E. Dieterle et al. 2017a). Briefly, fluorescent cells can be visualized using fluorescence microscopy only when proteins bind to the cell surface. Furthermore, the intensity of the fluorescence (that correlates to the amount of protein bound to the cell surface) can be quantified by flow cytometry.

The effect on CBM2 binding in H410F and H410F-D497L-D498L mutants was evaluated (**Figure 6**). In correlation with the results from the *in silico* analysis, the impaired binding of the H410F mutant was evidenced by a lower intensity of the fluorescence signal compared to the WT protein. (**Figure 6-A**). The H410F-D497L-D498L mutant could not bind to the cell surface and the signal observed was equal to the autofluorescence of *L. casei* BL23 cells alone. The findings were verified using fluorescence microscopy. As illustrated in Figure 4-B, the wild-type chimera (GFP-CBM2) was able to fully decorate bacteria In contrast, visual differences between the H410F mutant and the wild-type could not be observed, but fluorescence intensity was measured using cytometry assays. Unfortunately, bacteria decorated with the CBM2-triple mutant protein were not visible under the microscope, so only fluorescence intensity could be measured by cytometry.

**Figure 6:**
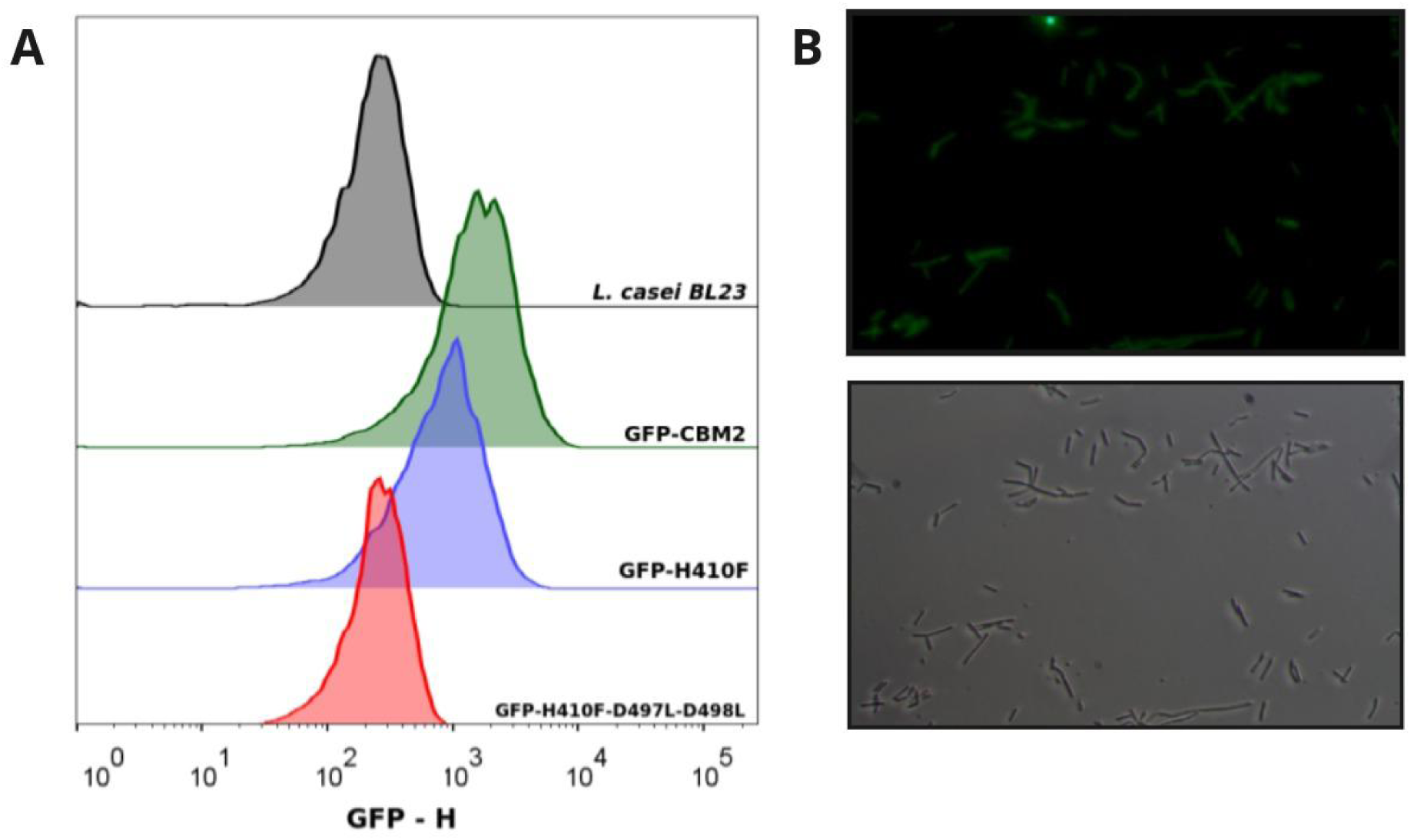
GFP-CBM2 binding assay to *L. casei* BL23. **A.** Analysis of GFP-CBM2 WT and mutant derivatives by flow cytometry. **B.** Analysis of GFP-CBM2-binding to *L-casei* BL23 by fluorescence microscopy.

In conclusion, these results highlight the significance of developing bioinformatics tools to generate accurate information and minimize the variables in the system, allowing us to focus on testing only the most crucial variables in vitro.

## Discussion

Many biological processes involve proteins that recognize carbohydrates. Some viruses use carbohydrate-binding proteins to recognize their host cells, bind, and inject their genetic material (Andres et al. 2010; Desmyter et al. 2013; Hayes et al. 2019; Fukuda 2012). There are experimental methods to determine which molecules are involved in virus-receptor recognition; however, the exact mechanism of the carbohydrate-binding process remains unclear (Gimeno et al. 2020). Understanding these interactions at the atomic and molecular level is crucial in the rational design of drugs that can prevent virus adsorption, and computational methods can be useful in addressing this challenge.

*Lactobacillus casei* J-1 phage recognizes its hosts through the baseplate protein machinery where the CBM2 domain mediates the interaction. It has been proposed that CBM2 recognizes cell wall polysaccharides in lactic acid bacteria that have Rhamnose in their structure, and this sugar was capable of inhibiting phage-host interaction (M. E. Dieterle et al. 2014a; M.-E. Dieterle et al. 2017a; Yokokura 1971). Moreover, the involvement of Rhamnose, present in CWPs of Gram-positive bacteria, in host recognition has been extended to other phages (Guérin, Kulakauskas, and Chapot-Chartier 2022). All this previous data reinforced the hypothesis that the CBM2 recognizes the CWPS during the phage adsorption step but the details of the interaction at the molecular level is still unknown.

In this work, we developed a pipeline to detect key amino acids responsible for the interaction between the CBS of CBMs and CWPs using the CBM2 domain of bacteriophage J-1 as a model. Our approach utilizes bioinformatics tools, including molecular docking and molecular dynamics, as well as freely available web servers to enhance the analysis process. The docking algorithms allow predicting with a reasonable degree of confidence how the ligand-protein complexes are formed, but they are not very efficient for carbohydrates, mainly oligosaccharides (Xiong et al. 2015; Perez and Makshakova 2022). Many strategies have been proposed to solve this, such as the SLICKfunction for BALLDock or the CHI energy function in Vina Carb, but many of these have disadvantages (E. D. Boittier et al. 2020; Kerzmann et al. 2008). To circumvent these drawbacks, we use the solvent structure information around the corresponding binding site to perform a *bias docking method (Arcon et al. 2021)*. The variability of the receptor was an important variant to consider for obtaining improved docking results. Furthermore, to reduce ligands complexity we also explored the probable conformations in solution of the different variants of CWPS using *explicit solvent MD* discarding unlike conformations. The combination of different methods allowed us to determine patterns of interaction. Residues H410, N412, Q586, and D498 were the most relevant in the protein-ligand interaction and, as expected, they tended to bind mainly to Rhamnose (Figure 4, panels C and D). Furthermore, we discovered that molecular dynamics can uncover residues not detected by docking. Specifically, we observed that the most common hydrogen bond interactions were established through the residues D498 and D497. By performing *in silico* mutations on the key amino acids, we determined that among all the generated variants, molecular dynamics free energy is an effective method to measure the loss of interactions and determine which variants are most sensitive. Our interaction energy delta calculations revealed that the presence of the H410F variant significantly decreases ligand interaction, highlighting its crucial role in binding (as shown in Figure 5 panels B and C). Furthermore, we assessed the affinity of the GFP-CBM2 constructs for the bacterial surface, including two proposed mutants, H410F, and a triple mutant H410F-D497L-D498L. The single residue H410F mutant still had the ability to bind to the cell surface but with reduced affinity compared to the wild-type protein. However, the H410F-D497L-D498L mutant completely lost its ability to bind to the cell, regardless of the protein concentration used (Figure 6), supporting the results from our simulation experiments.

Here, using the CBM2 domain as a model, we have demonstrated the efficacy of our proposed bioinformatics pipeline in identifying these crucial amino acids involved in carbohydrate recognition. To the best of our knowledge, this is the first established method to identify key amino acids in protein-carbohydrate interactions.

## Methods

### Computational Methods

#### Bioinformatic Analysis

For the search, alignment, and analysis of sequences, BLAST (https://blast.ncbi.nlm.nih.gov/Blast.cgi) was used (Altschul et al. 1990), conducting the corresponding searches against the RefSeq (O’Leary et al. 2016) (https://www.ncbi.nlm.nih.gov/refseq/) and UniProt (https://www.uniprot.org/help/uniref) databases (The UniProt Consortium et al. 2022). Multiple alignments were performed using the freely available Clustal Omega program (http://www.clustal.org/omega/) (Sievers et al. 2011). The search for similar proteins and proteins with similar motifs was carried out using the advanced search tools of the Protein Data Bank (https://www.rcsb.org/). The identification of cavities was done using the FPocket program (https://fpocket.sourceforge.net/) (Sievers et al. 2011; Le Guilloux, Schmidtke, and Tuffery 2009), while the characterization of potential ligand-binding sites was performed using FTmap Server (https://ftmap.bu.edu/param/login.php?redir=/param/) (Kozakov et al. 2015).

#### Molecular Docking

As an initial receptor we utilized the Carbohydrate Binding Module 2 structure (CBM2) from J-1 Phage Distal protein (Dit). The protein was obtained from Protein Data Bank with PDB ID: 5LY8 (M.-E. Dieterle et al. 2017b). All the ligands used were built and minimized using the tool, Carbohydrate Builder, Glycoprotein Builder, Woods Group. (2005-2023) GLYCAM Web. Complex Carbohydrate Research Center, University of Georgia, Athens, GA. (http://glycam.org). Then ligands were minimized with Molecular Dynamics simulations too. Carbohydrates were constructed from the resolved structures for carbohydrates present in the cell wall of L. casei BL23 (Vinogradov et al. 2016b; Balzaretti et al. 2017). The receptor structure was prepared following the standard AutoDock protocol (Morris et al. 2009) using the *prepare_receptor4.py* script from AutoDock Tools. All non-polar hydrogens were merged, and Gasteiger charges and atom types were added. The ligand PDBQT was prepared using the *prepare_ligand4.py* script available in AutoDock Tools and all ligands conformations were kept rigid for docking experiments. The grid size and position were chosen to include the whole ligand-binding site resulting in a 90×90×90 grid box, considering the W440 residue detected by the bioinformatic analysis carried out in the first stage of the work as the center of the binding site (see Bioinformatic Analysis section). The spacing between grid points was set at 0.375 Å. AutoDock Bias protocol (Arcon, Modenutti, Avendano, et al. 2019) was applied to perform a biased docking experiment taking into consideration the water sites information. Briefly, the carbohydrates -OH groups tend to mimic the interactions of certain water molecules around the binding site. For this we use the positions of the “water sites” calculated by molecular dynamics to modify the respective energy grids (OA and NA maps) using *prepare_bias.py* script. This method has been tested and validated in previous works (Arcon, Defelipe, et al. 2019; Arcon et al. 2017b).

For each system, 100 different docking runs were performed and the results were clustered according to the ligand heavy atom RMSD using a cut-off of 2 Å. The Lamarckian Genetic Algorithm (LGA) parameters for each conformational search run were kept at their default values (150 for initial population size, 1×10^7^ as the maximum number of energy evaluations, and 2.7×10^4^ as the maximum number of generations). The docking results were further analyzed by visual inspection.

In this work we considered both results, having negative ΔE and those high population poses. The best poses selected were used to do the binding interaction analysis and the corresponding complexes were used for the Molecular Dynamics simulations.

#### Molecular Dynamics Simulations

For the Molecular Dynamics (MD) simulations, we used with minor modifications the protocol described by Modenutti et al (C. P. Modenutti et al. 2021). We utilized as apo (without ligand) structure the Carbohydrate Binding Module 2 (CBM2) from J-1 Phage Distal protein (Dit) (PDBID: 5LY8) (M.-E. Dieterle et al. 2017b). The same receptor was used for the in silico mutations and their corresponding MD simulations (see below). The CBM2-Carbohydrate complex used was the result of the docking experiments described in the previous section.

Starting from each structure (wild type and mutant), we prepared the systems and performed three replicas of a 100 ns all-atom MD simulation, using the AMBER package (Case et al. 2022). The systems were prepared with the tleap module from the AMBER package, using the ff19SB/OPC force fields for amino acid/water molecules, respectively (Tian et al. 2020). Standard protonation states were assigned to titratable residues (Asp and Glu are negatively charged; Lys and Arg are positively charged). Histidine protonation was assigned by favoring for each residue the formation of hydrogen bonds. The complete protonated systems were then solvated by a truncated cubic box of OPC waters, ensuring that the distance between the biomolecule surface and the box limit was at least 10 Å.

Each system was first optimised using a conjugate gradient algorithm for 5,000 steps, followed by 150 ps-long constant volume MD equilibration, in which the first 100 ps were used to gradually raise the temperature of the system from 0 to 300 K. The heating was followed by a 300 ps-long constant temperature and constant pressure MD simulation, to equilibrate the system density. During these temperature- and density-equilibration processes, the protein C_α_ atoms were constrained by a 5 kcal/mol/Å force constant, using a harmonic potential centred at each atom’s starting position. Next, a second equilibration MD of 500 ps was performed, in which the integration step was increased to 2 fs and the force constant for restrained C_α_s was decreased to 2 kcal/mol/Å, followed by 10 ns-long MD simulation with no constraints. Finally, a 100ns-long MD simulation was carried out, with no constraints and the ’Hydrogen Mass Repartition’ method, which allows an integration step of 4 fs (Hopkins et al. 2015).

All simulations were performed using the pmemd.cuda algorithm from the AMBER package (Salomon-Ferrer et al. 2013). Pressure and temperature were kept constant using the Monte-Carlo barostat and Langevin thermostats, respectively, using default coupling parameters. All simulations were performed with a 10 Å cutoff for nonbonded interactions, and periodic boundary conditions, using the Particle Mesh Ewald summation method for long-range electrostatic interactions. The SHAKE algorithm was applied to all hydrogen-containing bonds in all simulations with an integration step equal or higher than 2 fs. All trajectory processing and parameters calculations were performed with the CPPTRAJ (Roe and Cheatham 2013) module of the AMBER package. Images of the molecules were prepared using the Visual Molecular Dynamics (VMD) program (Humphrey, Dalke, and Schulten 1996). The plots were created using the Matplotlib Python library (Hunter 2007).

Simulations of the CBM2-carbohydrate complexes resulting from the docking experiments were refined by short 10 ns simulations, following the protocol tested by Makeneki et al (Makeneni, Thieker, and Woods 2018). This protocol allows to increase those most probable poses by discarding the poses that are not stable or are energetically unfavorable.

#### Solvent structure analysis

Starting from the crystal structure for apo-protein and protein-ligand complexes, the structures were refined using molecular dynamics simulations. Briefly, Molecular Dynamics derived Water Sites (MD-WS) are derived from explicit water MD simulations (Gauto et al. 2009). The position of the oxygen atoms of water molecules in successive snapshots are clustered together when they are closer than a user-defined cut-off (usually 1.4Å), and their center of mass defines the corresponding WS coordinates. Subsequently, the probability of finding a water molecule inside a volume of 1Å centered on the WS coordinates is computed with respect to the bulk solvent. When this value is larger than 2, the WS is kept for biased docking.

#### *In silico* mutagenesis and impact over protein-ligand evaluation

The generation and evaluation of the in silico mutants were performed with FoldX and the destabilization of the binding of the ligand was performed using the structure package from the pyFodlX library (L. G. Radusky and Serrano 2022) which allows the inclusion of ligands during the estimation of the difference in free energy (ΔΔG) due to the substitution of a particular amino acid.

For protein-ligand evaluation, calculations of root-mean-square deviation (RMSD), hydrogen bonds (HBond), and Molecular Mechanics Poisson–Boltzmann Surface Area (MMPBSA) were performed. RMSD was calculated from the DM simulations compared to the initial docking results structure, using Visual Molecular Dynamics (VMD) (Humphrey, Dalke, and Schulten 1996). HBond was calculated using the ’cpptraj’ module available in the AmberTools package. MMPBSA, on the other hand, was performed using ’MMPBSA.py’ module available in the AmberTools package. The refinement and selection of the most probable interactions followed the same methodology as the refinement of the complex results obtained from the docking tests, as explained in the Molecular Dynamics section.

#### Data analysis and image generation

For the analysis of pockets, sites, and motifs at the structural level, Visual Molecular Dynamics (VMD) was used. Images of the molecules were prepared using the Visual Molecular Dynamics (VMD) program (Humphrey, Dalke, and Schulten 1996). For the analysis and graphical representation of the frequency of interactions, python 3.10 libraries were used (Humphrey, Dalke, and Schulten 1996; McKinney 2022; Sievert 2020).

### Experimental Methods

#### Strains, bacteriophages, and growth conditions

*Lactobacillus casei* BL23 was grown in MRS medium (Difco, USA) at 37 °C under static conditions. Escherichia coli TOP 10 (Invitrogen, Carlsbad, EEUU) was used for cloning, and *E. coli* T7 (NEB, Ipswich, EEUU) was used for protein expression. *E. coli* strains were grown in LB broth (Difco, USA) at 37 °C under moderate shaking. When appropriate, antibiotics were added at the following concentrations: kanamycin, 30 mg/ml (Sigma, USA); chloramphenicol, 34 mg/ml (Sigma, USA).

#### Cloning procedures

The cbm2 region was amplified using *Lactobacillus* phage J-1 genome (Accession number: KC171646) as the template with the following primers: ED98 5’-agcaggatccCCCGATGAGACTGATGGTT-3’ and ED99 5-’tgaacgagctcttaGTCTTTGAAGCGGTTAGG-3’ (Dieterle et al 2017). The amplicon was cloned into the pET28-GFP (M. E. Dieterle et al. 2014a) by using BamHI/SacI enzymes (CBM2) and T4 DNA ligase (Promega, USA). The plasmid was named GFP-CBM2.

The directed mutagenesis to obtain the H410F and the H410F-D497L-D498L derivatives was done by GenScript (https://www.genscript.com) on the GFP-CBM2 plasmid. The presence of the correct mutations was corroborated by sequencing.

#### Protein expression

Recombinant plasmids were transformed in *E. coli* T7 (NEB, Ipswich, EEUU) strains. Cells were grown at 37 °C in LB medium until the OD_600nm_ reached 0.6, and protein expression was induced with 0.5 mM IPTG overnight at 18 °C. The cells were harvested by centrifugation (4000g for 10 min) and the pellet was homogenized and frozen in lysis buffer [50 mM Tris pH 8.0, 300 mM NaCl, 10 mM imidazole, 0.1 mg ml ^-1^ lysozyme, 1 mM phenylmethylsulfonyl fluoride (PMSF)]. After thawing, DNAse I (20 mg ml ^-1^) and 1 mM MgSO_4_ were added and the cells were lysed by sonication. The pellet and soluble fractions were separated by centrifugation (16000g for 30 min). Purification was performed on an AKTA FPLC system following an immobilized metal ion-affinity chromatography using a 5 ml HisTrap Crude (GE Healthcare) Niquel chelating column equilibrated in buffer A (50 mM Tris pH 8.0, 300 mM NaCl, 10 mM imidazole). The proteins were eluted with buffer B (buffer A supplemented with 250 mM imidazole).

#### Fluorescence binding assay

Cell binding assays using purified GFP-CBM2 (wild-type and mutant proteins) domains were carried out as described by Dieterle et al. 2017, with some modifications. Briefly, 300 μl of exponentially growing bacterial cells (*Lactobacillus casei* BL23 OD_600nm_ of 1) were centrifuged and resuspended in 300 μl of modified phage buffer (MPB) (50 mM Tris-HCl, 100 mM NaCl, 0.1% Tween 20, 10 mM CaCl_2_) and incubated with the indicated concentration of protein fusions for 20 min at room temperature. Cells were washed three times with phosphate-buffered saline (PBS) buffer, and binding to the bacterial cells was detected by fluorescence microscopy (Axiostar Plus; Carl Zeiss) with a 100X objective with oil immersion and phase contrast.

#### Flow cytometry

The same protocol described above for fluorescence binding assays was used and cells were analysed by BD FACSAria II flow cytometer to a 488-nm light source. The bacterial populations first were gated based on their SSC and FSC profiles and were recognized by sampling a blank PBS solution in parallel. Ten thousand events were recorded in each experiment. Fluorescent and nonfluorescent cells within the gated population were discriminated based on fluorescent intensity (GFP-H). BD FACSAria software (BD FACSDiva, firmware version 6.1.3) was used for data acquisition, and FlowJo v10 software was used for subsequent analysis.

